# Progress in Developing a Bark Beetle Identification Tool

**DOI:** 10.1101/2024.09.08.611906

**Authors:** G. Christopher Marais, Isabelle C. Stratton, Andrew J. Johnson, Jiri Hulcr

## Abstract

This study presents an initial model for bark beetle identification, serving as a foundational step toward developing a fully functional and practical identification tool. Bark beetles are known for extensive damage to forests globally, as well as for uniform and homoplastic morphology which poses identification challenges. Utilizing a MaxViT-based deep learning backbone which utilizes blocked local and dilated global attention to classify bark beetles down to the genus level from images containing multiple beetles. The methodology involves a comprehensive process of image collection, preparation, and model training, leveraging pre-classified beetle species to ensure accuracy and reliability. The model’s F1 score estimates of 0.99 and 1.0 indicates a strong ability to accurately classify genera in the collected data, including those previously unknown to the model. This makes it a valuable first step towards building a tool for applications in forest management and ecological research. While the current model distinguishes among 12 genera, further refinement and additional data will be necessary to achieve reliable species-level identification, which is particularly important for detecting new invasive species. Despite the controlled conditions of image collection and potential challenges in real-world application, this study provides the first model capable of identifying the bark beetle genera, and by far the largest training set of images for any comparable insect group. We also designed a function that reports if a species appears to be unknown. Further research is suggested to enhance the model’s generalization capabilities and scalability, emphasizing the integration of advanced machine learning techniques for improved species classification and the detection of invasive or undescribed species.

## Introduction

Computer vision technologies for the purposes of classification and identification are becoming progressively more prevalent in ecological research, biological systematics, and biosecurity. This is mainly attributable to the application of deep learning techniques, with convolutional neural networks (CNNs) and transformer networks being particularly influential (1,2). These methodologies have facilitated the emergence of hybrid architectures like MaxViT, which integrates the strengths of both CNNs and transformers to improve performance and efficiency, especially in contexts characterized by constrained data resources (3).

In ecological studies, deep learning models have demonstrated potential by accurately identifying species that are visually similar. These models excel in consistent, reliable, and accurate classifications even among morphologically similar species (4–9). This capability underscores the technology’s potential in ecology, biosystematics, and invasive species detection where precise species identification is crucial (4–9).

Bark beetles, an abundant and widespread group of wood boring insects, are notable for their potential to cause significant tree mortality. Their outbreaks are increasing particularly in scenarios where non-native species invade a region with naive hosts, or when the tree hosts are stressed and lessen their defense abilities, often because of climate change (10) The importance of detecting and accurately identifying these beetles cannot be overstated for forestry industries; however, existing identification methods fall short in efficiency and scale, posing challenges in safeguarding forests worldwide against infestations(11).

Bark and ambrosia beetles, encompassing Scolytinae and Platypodinae within Curculionidae, represent a diverse group with over 8000 species spanning approximately 250 genera across 26 different tribes (12). Although most bark beetle species are not typically categorized as pests, certain species can cause significant forest damage (13). This uneven taxonomic distribution of pestiferous species makes species-level identification critical: some species can trigger a region-wide regulatory action, while others with a very similar morphology are entirely harmless.

Global trade and travel have facilitated the spread of invasive species, enhancing the risk posed by these beetles (14).

The ecological impact of bark beetles is significant, as they alter habitats and affect numerous organisms (15). In natural ecosystems, bark beetles are important ecosystem engineers (reference), but in managed stands and plantations, some species cause a notable economic damage (16). The economic damage in industries like pulp, paper, and timber, with the southern pine beetle alone causing an estimated annual loss of around $43 million in the U.S. South (17). The increasing prevalence of bark and ambrosia beetles, exacerbated by climate change and international trade, highlights the urgent need for effective management strategies.

The monitoring, detection and identification of current and future bark beetle pests poses a significant challenge due to their small size, morphological homogeneity (18–20), and often large volumes of material in traps or collections. Traditional methods of identification, such as visual classification, require considerable expertise and are time-consuming. Molecular methods, like DNA sequencing, are impractical for large-scale pest detection efforts due to their slow processing, high costs, and technical issues (21).

Currently, federal, and state-level programs like the Cooperative Agricultural Pest Survey (About the CAPS Program | CAPS, 2023) and the USDA Forest Service’s Early Detection and Rapid Response program (11) rely on human expertise to monitor and identify invasive pests. While trained identifiers are generally skilled, the sheer volume of specimens can hamper the identification process, by inducing errors and focus fatigue. Moreover, the logistical challenges of curation of the voluminous non-target bycatch limits the development of physical as well as digital reference collections. Integrating artificial intelligence tools could help human experts more efficiently identify uncommon or unknown specimens, thus improving identification speed, refining accuracy, and preserving valuable reference material (23).

This study tests a deep learning model capable of classifying bark beetles to the genus level from images of multiple beetles at a time. Although we focus on genus-level classification here, achieving species-level identification remains the ultimate goal— especially for identifying and managing new invasive species. We also recognize that bark beetle taxonomy is not static. Ongoing revisions and reclassifications mean that species definitions and genus and tribe placements change with additional data. Future iterations of our approach can incorporate these taxonomic updates, retraining the model as needed to maintain or improve its accuracy, thereby ensuring that the tool remains robust and useful in the face of evolving species definitions.

We specifically focused on the development of a precursor model that could be used and expanded for practical, existing identification applications which typically deal with multi-species and multi-specimen samples. We tested a model structure that first disaggregates compound photographs, and then identifies individual beetle specimens. To train the model, we also devised a method for efficiently generating aggregate photos of multiple individuals, yielding the largest bark beetle photo collection ever assembled. In addition, we also devised a method for determining a probability that a genus is “unknown” (absent in the training set), which is important in detecting novel genera or species.

## Methods

Images were collected by photographing physical bark beetle samples that have previously been classified to species by bark beetle taxonomists according to current classification (24). We included only one species per genus. The selection of the beetle genera for the model training was based on three factors. Firstly, their economic importance was a key consideration, ensuring inclusion of beetles with significant impact in industries and ecosystems. Secondly, accessibility of large series of physical samples was required. Lastly, consideration was given to the prevalence of these genera in traps of the national pest monitoring networks, where the model may be helpful in the future.

### Training set generation

The 12 selected genera of pre-classified bark beetles were stored in 70% ethanol at −80°C before and after photography. During image collection the samples were placed in petri dish on a white background and submerged in 70% ethanol. Multiple bark beetles were included in a single image using a macro photography technique designed to optimize reproducibility of the image capturing process and image quality (Bark and Ambrosia Beetle Macro Photography for an AI Training Dataset, 2024). This approach to capturing images of multiple beetles at once sped up image collection compared to capturing one image of a beetle at a time. Each batch of beetle samples was of the same species to ease digital labelling. Each batch of beetle samples were also photographed 10 times over after agitating the peri dish between photographs. This was to increase the number of visual angles for each individual beetle photographed.

Pictures were taken with a Canon EOS REBEL with a 60mm lens. We kept the ISO at 100 and used a ring flash to reduce the shadows around each beetle. Each image taken in this way resulted in an image of multiple beetles at a resolution of 3456 x 5184 with 72 dpi at 24-bit color depth. The images with multiple beetles in it will be referred to as a composite image. The intention was to keep the photography setup relatively straightforward to enhance reproducibility in the future. All images were taken from the top down onto the beetles on a rig that held the camera. The white background was selected to control the background with a high contrast between the beetles to aid in data preparation and identifying individual beetles.

### Data Preparation

The composite images of multiple beetle samples were split into multiple images. Resulting in multiple images with one beetle sample in each image. This was done using the Scikit-Image python package. This task was not a part of the deep learning model and kept as a separate process in the pipeline. We did this because our focus remained on optimizing the bark and ambrosia beetle classification problem and not general object detection.

The first step in data preparation was to confirm that all images were in focus and had enough lighting. The composite images were split into test validation and training datasets. The test set was 20% of all the composite images, maintaining the ratio of genera in the training data to avoid skewing the testing or training sample in that way (Figure 1). Thereafter the remaining 80% of the training data were split using 5-fold cross validation (26). This results in five replicates of the data to monitor for sampling biases. The image data was split along the composite images and replicates of each composite image to avoid cases where the same individual may be in both the testing (or validation) and training dataset.

**Fig 1:**
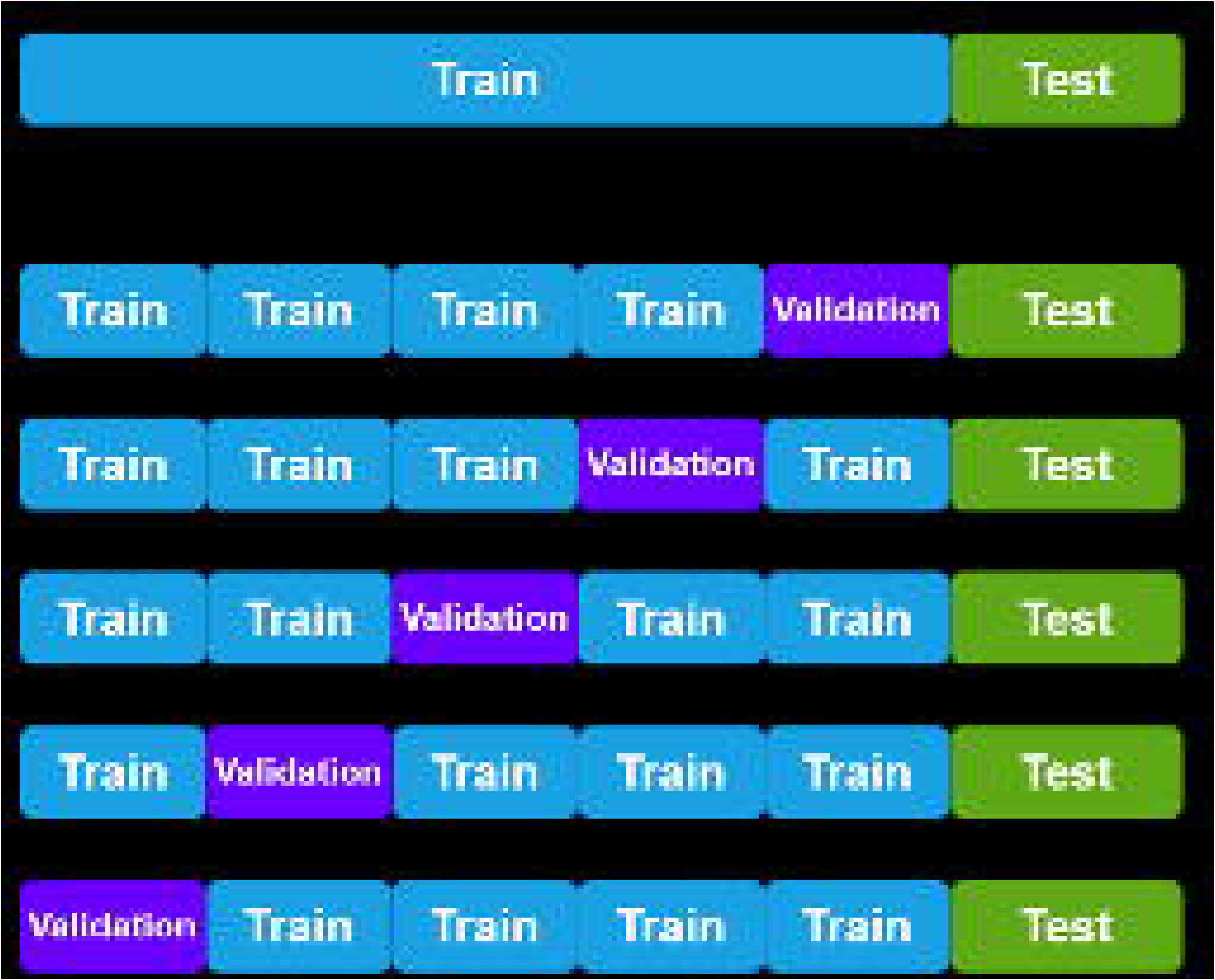
Example of 5-fold Cross-validation.

Once the image data were split, the composite images were broken down into smaller images each containing a single beetle (Figure 2). This was done using the Scikit-Image package in python using the following steps (27).

1. Import color composite image.
2. Convert image to black and white.
3. Binarize the image with beetles as black and the background as white using the Otsu thresholding algorithm (28).
4. Remove any black objects that are in contact with the image edge.
5. Detect all black blobs in the image and draw rectangle boxes around each of them using the built-in label function from Scikit-Image.
6. Use K-means with 2 clusters separating objects by size
7. Keep only the larger objects.
8. Use the rectangle coordinates of kept objects and extract those sections from the original color image.

**Fig 2:**
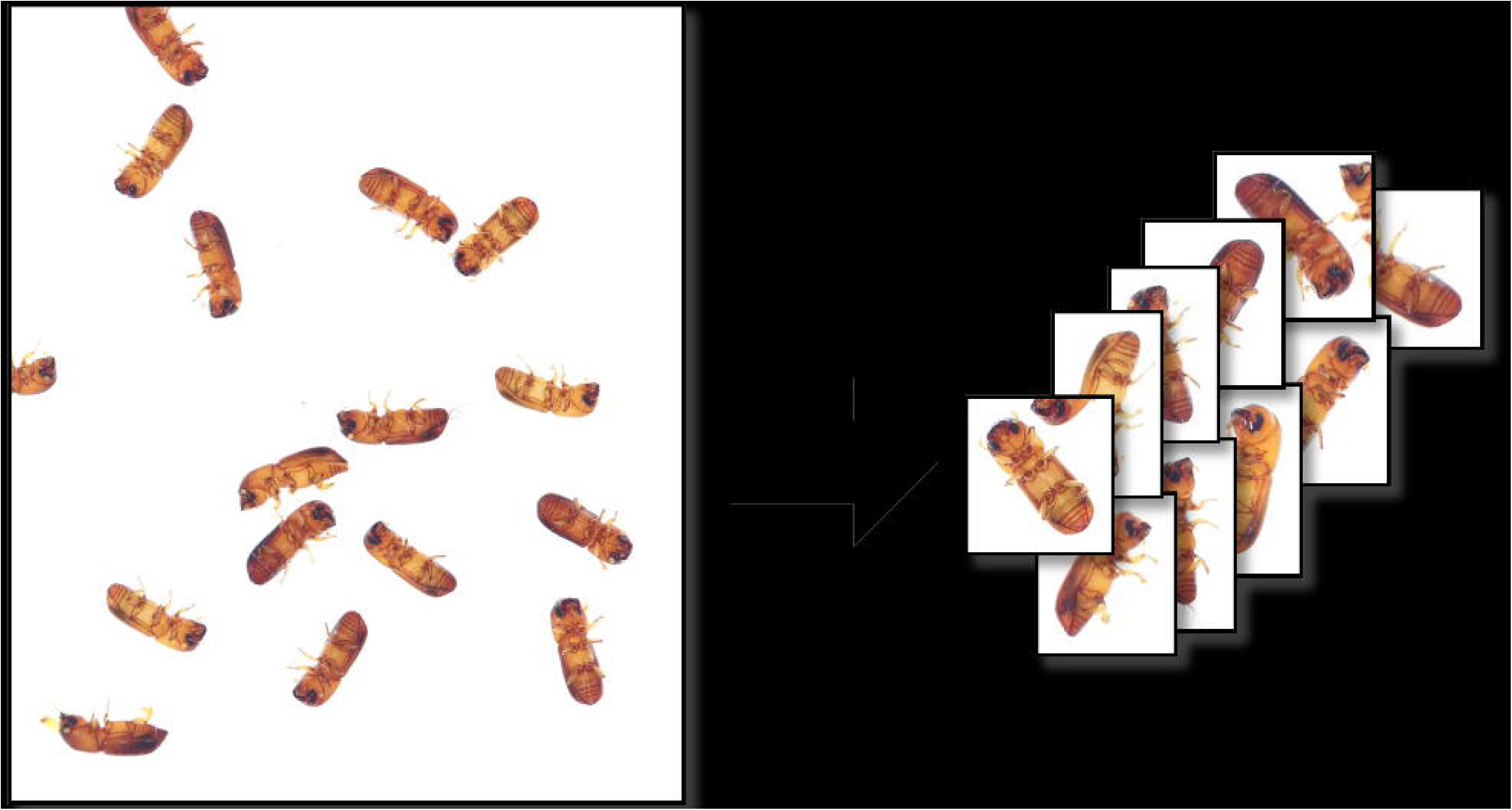
Splitting of Composite Image to Individual bark Beetle Images

Images that contained multiple beetles or contained only partial segments of beetles were manually removed from the data through visual inspection of all files. Images that contained any foreign bodies or organisms were also removed.

### Model Training

We adopted a transfer learning approach for model training to reduce computation time and amount of data required to train the model effectively (29). We utilized a MaxViT architecture that has 116.1 million parameters and 92.6 million activations, previously trained on a subset of the ImageNet-22K dataset, comprising over 14 million annotated images and 22 thousand labels (3,30). The pre-trained model was obtained through the Pytorch Image Models (timm) package on Hugging Face (31). Specifically, it was trained on the ImageNet-12K subset, consisting of 1.2 million images across 11,821 classes, and thereafter it was further fine-tuned on the ImageNet-1K subset, containing 1.2 million images with 1000 classes.

We opted for the MaxViT model architecture to leverage its blend of global and local attention mechanisms (3), predominantly employing the Adam optimizer (32). The model utilizes various activation functions like ReLU, SiLU, and GELU (3,33), with Tanh being crucial in the linear classification head. In the classification layer, the model processes the feature map into a vector of 75,264 values, which is then reduced to a vector of length corresponding to the number of classes (12 in our case, corresponding to the number of beetle genera). The Tanh function produces values between −1 and 1, which are converted into prediction probabilities using SoftMax (34). This ensures that the output probabilities sum up to 1 for all classes. The consistent label smoothing cross-entropy loss function was applied to estimate the error (loss) for all models (35), this specific objective function was chosen for its suitability in multi-class classification tasks like ours. This loss function also serves as a regularization method, mitigating overfitting and enhancing model performance and generalization by introducing slight label noise to the training data (36).

The pretrained and fine-tuned MaxViT model was further trained on our images that had dimensions of 224 x 224 pixels to suit the resolution that the model was pre-trained on. The conversion was carried out using the Pytorch and FastAI packages in python (37,38).

To artificially increase the size and diversity within the training data and to prevent overfitting to our data and improve generalization of the model we used image augmentations (Shorten & Khoshgoftaar, 2019). Each of these augmentations were applied to a random degree during the training process with a probability of 0.8 to be applied to any image in each batch. The augmentations we applied included gaussian blur and random patches of noise. Additionally, we performed changes to the rotation, flip, brightness, contrast, zoom, scaling, warping, and cropping of images as augmentations.

During training we replaced the existing classification head of the model with our own classification head with one node for each class (12 nodes in total). The model then went through a warm-up phase of one epoch of training where the layers of the model were all kept frozen excluding the classification head (40). Thereafter the model was trained for several additional epochs where no layers were frozen until validation loss of the model plateaued and converged with the training loss (Figure 3).

**Fig 3:**
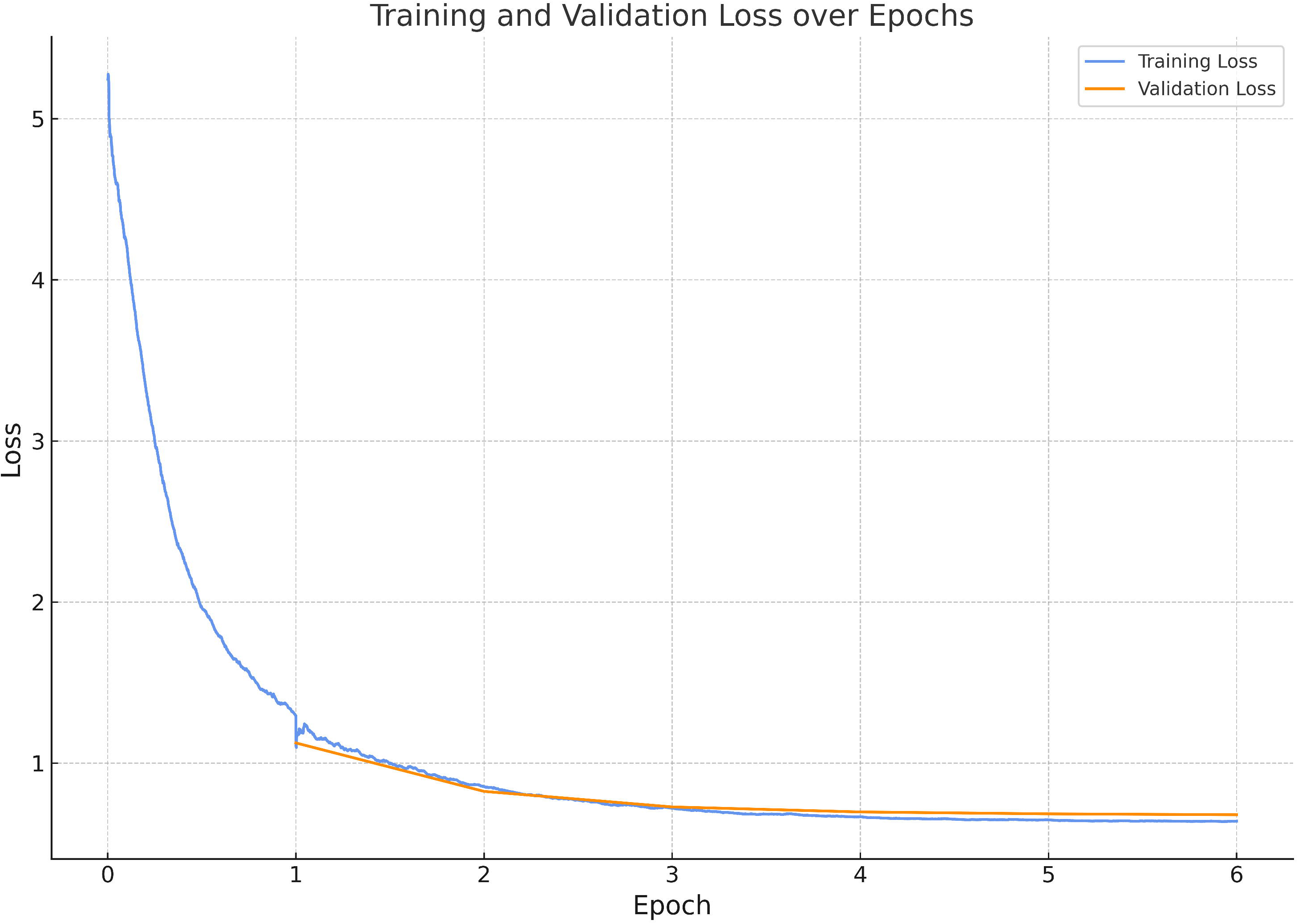
Training Loss (blue) and Validation Loss (orange) Across Epochs for Best Model During Parameter Tuning. The x-axis represents the number of epochs, and the y-axis represents the calculated loss.

The next step was to identify the ideal set of hyperparameters to train our model with. To do this we made use of the Hipergator supercomputer at the University of Florida. During hyperparameter tuning, we utilized PyTorch and FastAI for GPU-accelerated model training, employing the Weights and Biases (WandB) package for overseeing parameter optimization (41). Bayesian search was employed to expedite parameter optimization instead of exhaustive grid search. The Hyperparameters investigated included batch size, epochs (excluding the warm-up phase epoch), and learning rate. These were selected based on their impact on the validation loss. After evaluating various models, a batch size of 64, trained for 5 epochs with a learning rate of 0.003, was chosen. These hyperparameters were subsequently utilized for all experiments and training of the parameters for the final model.

For each evaluated image, the model generates probabilities for each of the 12 classes, corresponding to specific beetle genera. However, it lacks the ability to classify unknown genera. Training a class of entities unknown to the model would require impractical amounts of training data. A common alternative for detecting unknown classes is to apply a decision threshold on the existing probabilities which determines whether an output is classified as unknown or known. However, this approach does not produce a probability of how confident the model is that the class is unknown (42).

To address this limitation, we propose a formula to estimate the probability of the unknown class based on the distribution of probabilities of trained classes, aiming to provide users with additional information for decision-making during deployment.

Let *P* be a vector of length *n* representing the probabilities of *n* classes. Let *t* be a scalar representing the value of the threshold we use in our algorithm to represent the decision boundary. We define the probability *P*_*u*_ as the unknown class or out-of-class probability (Equation 1). Where *d* is a vector of length *n* representing each probability *P* to the power of the compliment of the absolute difference between each of the probabilities *P* and the threshold *t* (Equation 2).

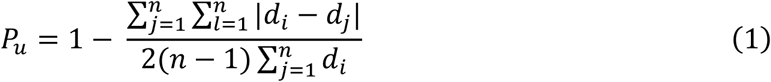

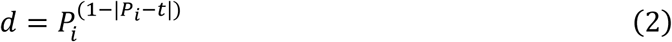

This formula generates an estimate probability that an image belongs to the unknown class as a function of how much higher a single probability in the vector is than the other probabilities and whether it is above the threshold parameter. The ideal vector has one high probability, above the threshold, and the rest of the probabilities are very low. This will produce a low *P*_*u*_ close to 0 unknown class probability estimate because the classifier seems to be certain that one class is the true class. However, if a vector is produced with multiple probabilities that are all very similar the unknown class probability will be closer to 1. This makes it possible to produce an estimate of the unknown probability and represent it in an output alongside the class probabilities as a measure of certainty of the classifier without sacrificing the ability to use a threshold on the original output of the model for out-of-class classification.

The code to using this model and a demonstrative version of the bark beetle image classifier are available for public access on Hugging Face (https://huggingface.co/spaces/ChristopherMarais/Andrew_Alpha).

## Results

The model achieved a high level of accuracy for the bark beetle genera included in this study, with an F1-score of 0.99 on the validation dataset and 1.0 on the testing dataset. While these metrics indicate excellent performance under controlled conditions, the model’s ability to distinguish closely related species or handle more diverse real-world scenarios remains to be assessed.

We accumulated 2 290 composite images of 12 genera (Figure 4). We were able to obtain an average of 12.9 single beetle images from each composite image (Figure 6). The average number of single beetle images per composite image differed markedly between genera (range: 1248-4968, Figure 5). This was because the beetles were all different sizes, and it wasn’t possible to fit the same number of beetles in each composite image.

**Fig 4:**
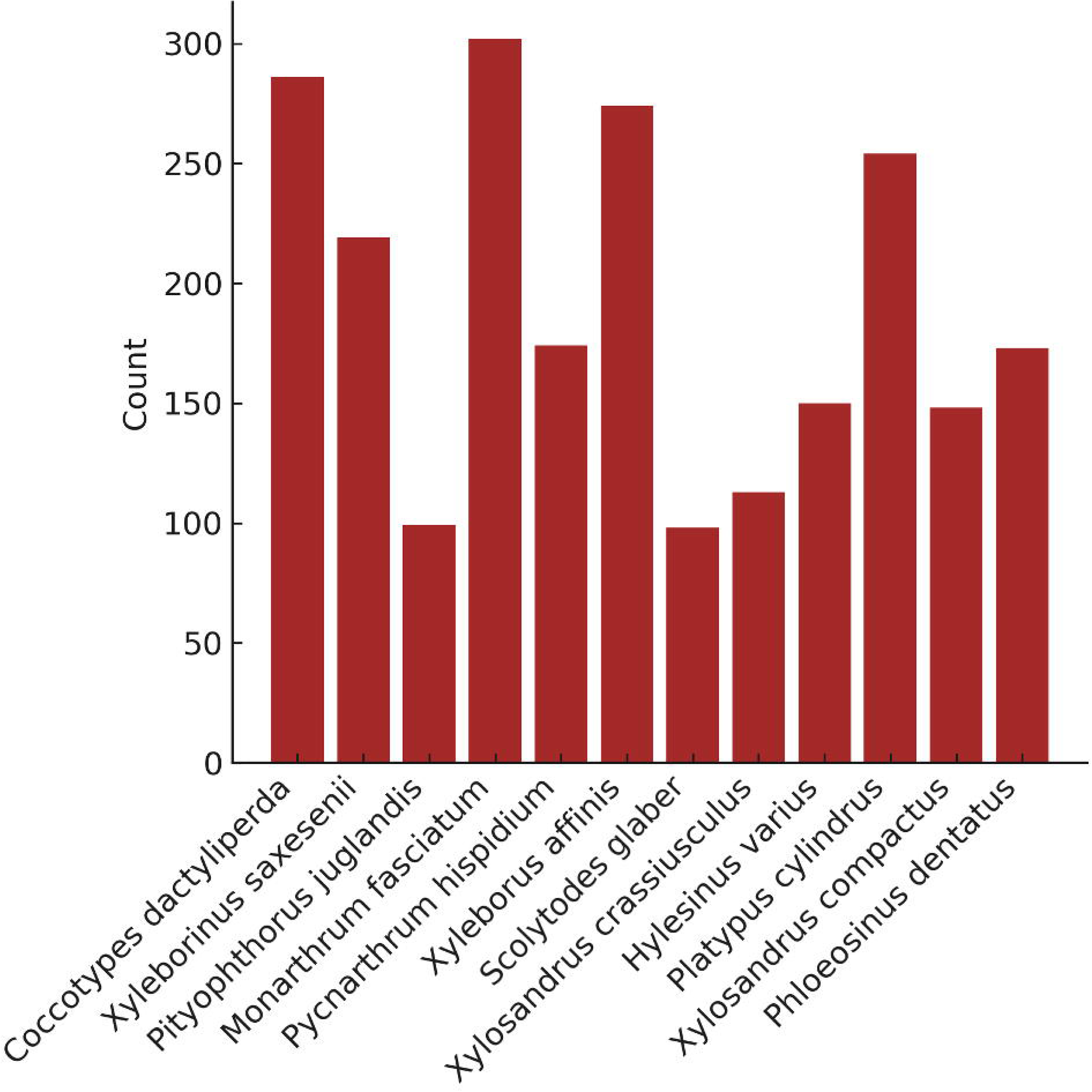
Number of Composite Images Collected per Genus

**Fig 5:**
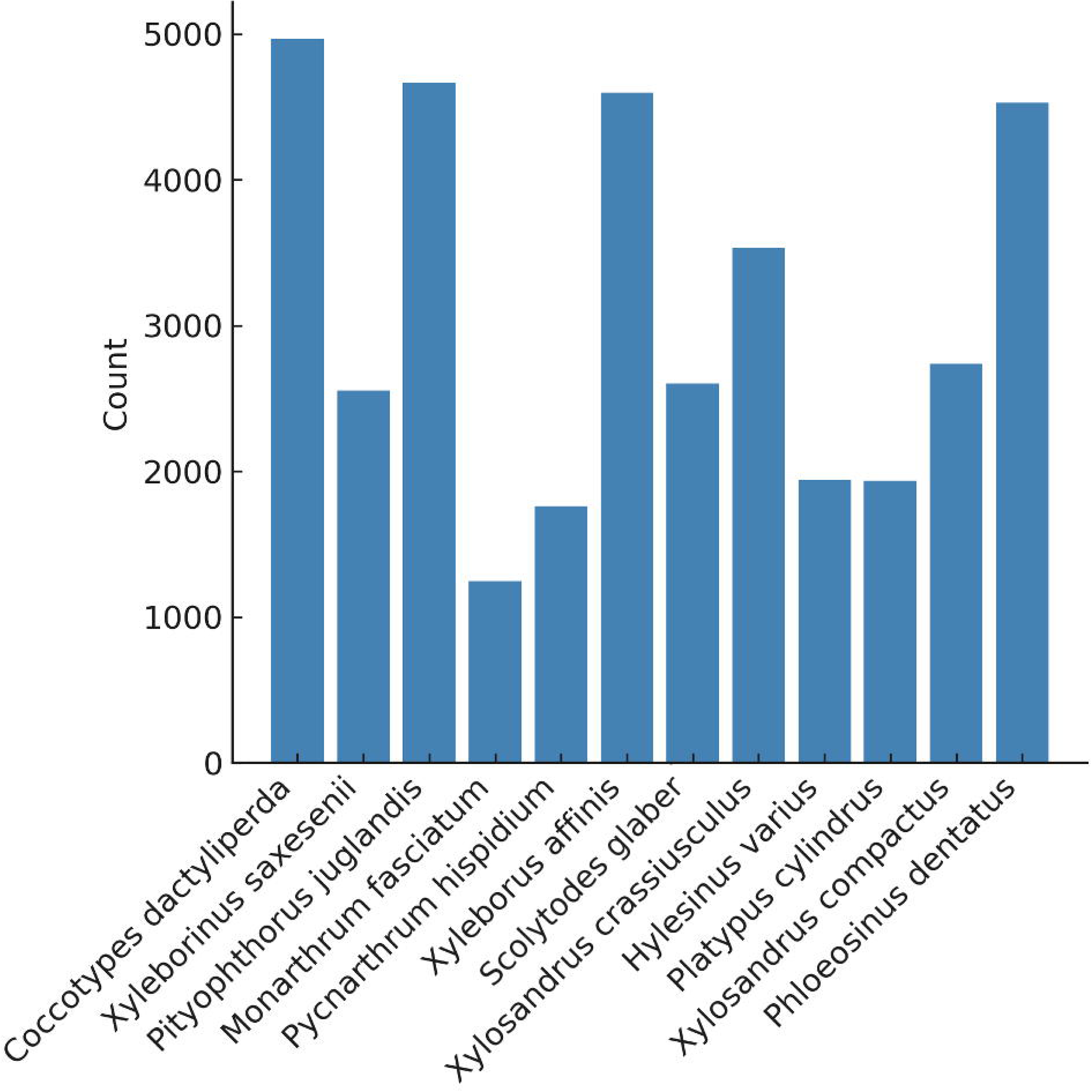
Number of Single Beetle Images Generated per Genus

**Fig 6:**
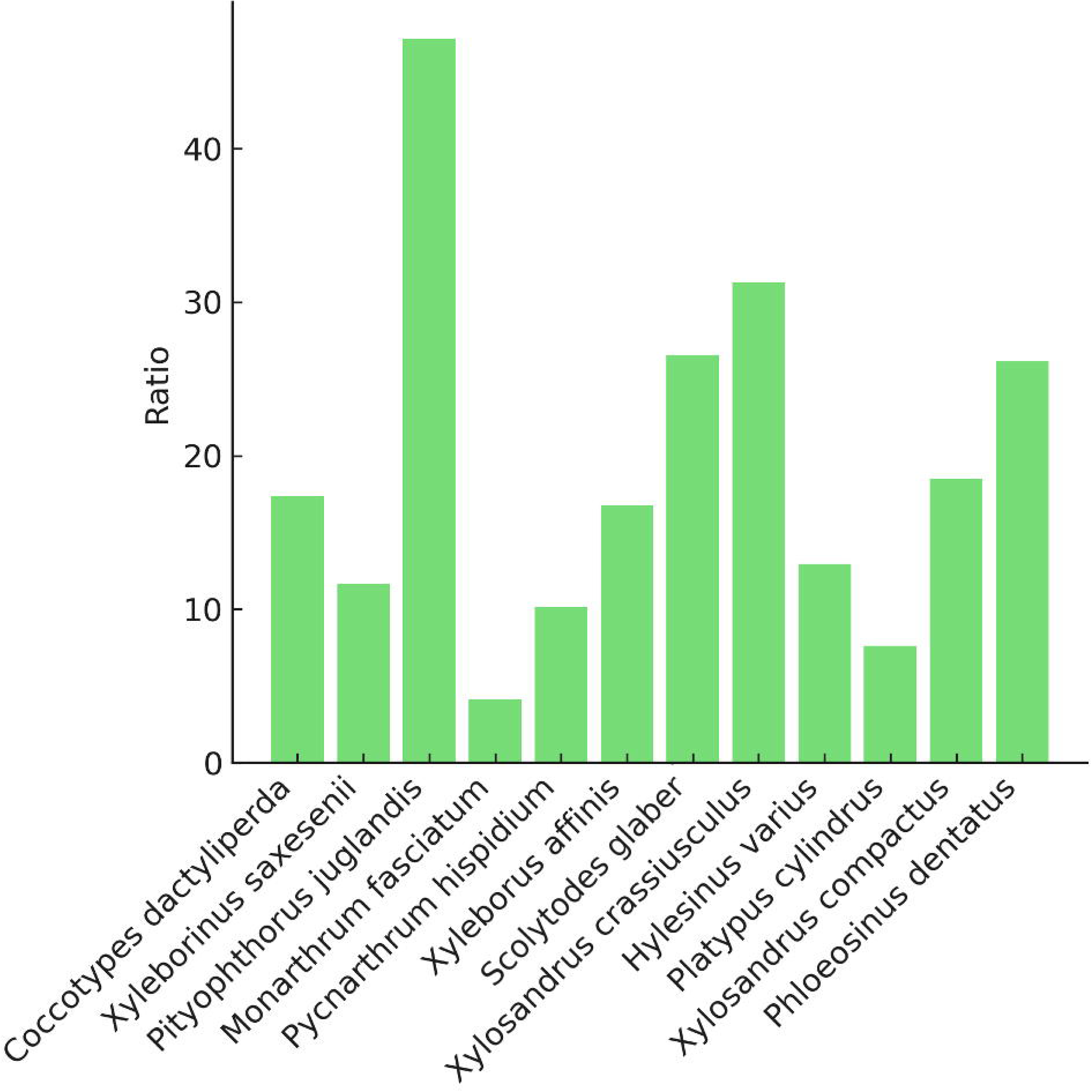
Average Number of Single Beetle Images per Composite Image by Genus

After settling on a batch size of 64 images with a pre-trained model, and 5 epochs with a learning rate of 0.003, these parameters were used for further training, evaluation, and testing. To evaluate the model thoroughly we used 5-fold cross validation. This meant that for every fold we evaluated the model a different subset of the data covering all the training data across all folds. For each fold the model was also evaluated on the holdout testing data. To evaluate the model, we made use of F1-score instead of a general accuracy score. The F1 score is more suited to multi-class classification as it is more sensitive to data distribution (43). An F1-score ranges between 0 and 1 with higher values indicating better classification. The F1-score obtained for the model averaged over all the folds for the validation data was 0.99 and for the testing data was 1.0.

To estimate our model’s performance on out-of-class classification with our unknown class probability formula we chose to test it on images that are closely related to the training data, but not included in the training data. This was done by training multiple models each with one class of beetles excluded from the training data. Thus 12 different models, each of which has been trained only on data of 11 classes and excluding one class. The excluded class was still used to evaluate and test the model as and represented a true label of “unknown”. This was repeated for each of the 5 folds to get a robust estimate of performance. Therefore, a total of 60 models were all trained using the originally optimized set of parameters. For each model we then applied the standard threshold decision boundary method of out of class classification to test the ability of the model to identify an unknown class. The same was repeated using our own unknown class probability formula to estimate the probability that an image belonged to an unknown class. Thereafter the class with the highest probability was selected as the classified class.

Both the conventional decision boundary approach and our unknown class formula approach add an additional threshold parameter to the inference process. This threshold parameter can be optimized by using the Receiver Operating Characteristic (ROC) curve. A ROC curve is a plot that shows the performance of a model at various threshold values. This plot compares the True Positive Rate (TPR) to the False Positive Rate (FPR). The TPR is a synonym for recall and is the ratio of true positives to the sum of true positives and false negatives. The FPR is the ratio of the false positives to the sum of false positives and true negatives. We calculated the FPR and TPR for the unseen testing data and the validation data for each model. We took the average of the ROC curves for each fold and compared them. We decided to take a One-vs-Rest approach on the ROC curve with the unknown class as the positive class and all other classes as the negative class(44).

The ROC and the conventional thresholding approach perform similarly both close to the ideal curve and far from the random curve (Figure 7). The ideal curve is in the top left corner with a TPR of 1 and a FPR of 0. The ideal area under the curve (AUC) is therefore 1. The conventional threshold decision boundary approach comes close with an AUC of 0.981 and our own formula approach is very similar with an AUC of 0.982. This means that both methods work well at classifying an unknown class. The optimal threshold can be obtained as the point on the plot that is closest to the 1 TPR and 0 FPR point. Both methods suggest an optimal threshold of 0.8 (Figure 7).

**Fig 7:**
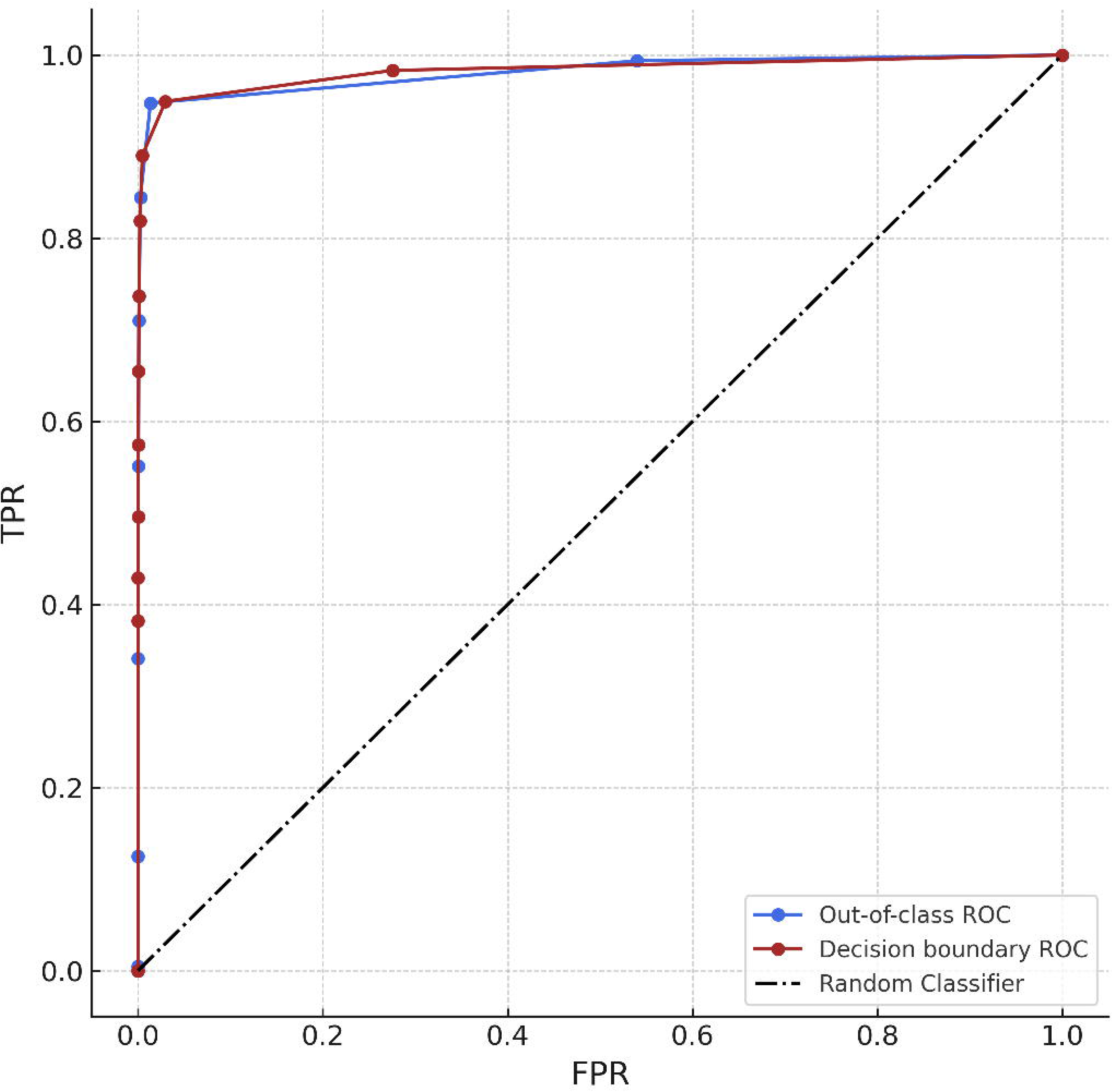
ROC curves of our out-of-class algorithm (blue) and a threshold decision boundary (red) representing the true positive rates (TPR) compared to the false positive rates (FPR) for test data classification results across all folds. The Diagonal (black) line represents the performance of a random binary classifier.

To get an estimate of how well the model performs in general when using this threshold, we calculated the weighted F1-score using the threshold. This gave us an F1-score of 0.97 for our formula and an F1-score of 0.98 for the conventional decision thresholding approach when applied to the unseen testing data.

## Discussion

In this study, we developed an advanced tool for identifying bark beetles, a significant threat to forestry due to their potential for extensive tree mortality. The need for efficient and accurate identification methods is underscored by the limitations of the current approach, which relies almost entirely on trained personnel, which is difficult to scale globally to the level of need. By leveraging a deep learning model, specifically a MaxViT architecture capable of classifying bark beetles down to the genus level from images, the study represents a significant leap forward. However, the current implementation differentiates bark beetle genera rather than achieving fine-grained species-level identification. Future work will target species-level resolution to improve invasive species detection.

The methodology involved collecting images of pre-classified beetle species, which were then used to train the model through a rigorous data preparation and model training process. This approach not only promises to enhance identification accuracy but also addresses the challenges posed by traditional identification methods. The model was evaluated using F1 scores and showed notable accuracy in identifying the 12 genera on which it was trained. In addition, its ability to classify unknown genera makes this approach even more promising for bark beetle detection and research.

These high identification scores indicate that the model can reliably classify the included genera in our controlled setting. However, additional testing is needed to confirm its precision and reliability under more variable and challenging conditions. Additionally, the model’s out-of-class classification approach shows promise for detecting previously unencountered genera, though further validation is required to confirm this potential in practical applications.

Several features of the model requires further testing before deployment in real-world scenarios with higher noise levels and greater variability in image data. Our training set collection and preparation involved the photography of beetle samples within a controlled environment, which may not faithfully replicate the diverse conditions encountered in natural settings. In the future it may be worth investigating the effect of applying this model to samples states and conditions that are common in pest research but have not been included in our training set (e.g., dry pinned beetles, beetles directly from traps, etc.). In the future, applying the model to images from different imaging devices may also be valuable in evaluating the true generalizability of the model.

The process of obtaining single beetle images from composite images, integral to preprocessing, was based on a rudimentary algorithm. The model’s use in scenarios with complex or varied backgrounds will require further development of the disaggregation step.

Our study also contended with data imbalance issues, characterized by unequal image distribution among various beetle species. In theory such imbalance could lead to a model bias, favoring more frequently represented species, over lesser-represented species. However, this does not seem to have affected accuracy with our initial set of 12 species. The high F1-scores, while indicative of the model’s performance, also raise the possibility of overfitting, particularly given the homogeneity of the training dataset. The methodology employed for out-of-class classification, focusing on unknown genera identification, was based on a custom formula that relied on the distribution probabilities of known classes.

Concerns regarding scalability and data efficiency of the model were also noted, particularly in relation to incorporating new species into the dataset. For a model intended to identify a broad array of species in diverse environments, efficient data collection and processing capabilities are imperative.

Additionally, reliance on pre-trained models, while beneficial in some respects, introduces the risk of inheriting biases from the source datasets (45).

Considering these limitations, we propose several directions for future research to enhance the model’s practicality and scientific utility. Expanding the number of species in the training set and improving data efficiency are crucial steps in expanding the model’s practical value. A significant focus should be placed on refining the object detection algorithm to improve accuracy, particularly for images with higher noise levels. Our current model distinguishes beetle taxa at the genus level. Achieving true species-level identification is critical for detecting emergent invasive species and will require hundreds more species in the training set, and refinement of the model’s classification head. The species selection should be near-exhaustive for a given fauna.

To enhance the model’s capabilities further, a multifaceted approach involving the integration of advanced machine learning techniques can be considered. A pivotal enhancement would be the integration of out-of-class classification directly into the network’s classification head. This integration is crucial as it would enable the model to identify and categorize species that were not part of the original training set, effectively recognizing ‘unknown’ or novel classes. By doing so, the model would not only classify known species with high accuracy but also flag species that are beyond its current knowledge base. This feature is important since in practice, encountering unclassified or new species is common, and in some application contexts, they ultimate goal.

In addressing the challenges posed by data gaps and underrepresented species, the incorporation of generative components into the model could be transformative. This approach would involve using generative techniques to synthesize training data for species that are not sufficiently represented in the existing dataset. This method could significantly enhance the model’s training process (46).

Another significant enhancement could be realized by adopting a self-supervised learning approach (47). By applying this approach, the model can learn from unlabeled data, which is a common challenge in ecological datasets where labeling can be resource-intensive and time-consuming. This approach enables the model to develop a more nuanced understanding of the features that distinguish one species from another, even in the absence of explicit labels. Finally, field deployment of the model warrants careful consideration of the needs of the research and regulatory community, ensuring that it is non-disruptive and user-friendly.

## Conclusion

A dual-pronged approach, combining advanced machine learning techniques with a comprehensive data management strategy and an image disaggregation algorithm, enabled us to classify morphologically similar insect genera from a computer vision perspective. Central to our methodology was the deployment of a MaxViT-based machine learning model, optimized for beetle image analysis. While we currently classify at the genus level, ongoing research aims to refine the model to reach species-level resolution. This enhancement is crucial for early detection of new invasive beetle species and more precise management decisions.

To augment the model’s utility and scientific contribution, future research should focus on expanding the training dataset to cover as large a proportion of the beetle species diversity as is practical, refining the object detection algorithm, incorporating contextually appropriate machine learning techniques that focus on addressing high class imbalances and generative components, and exploring self-supervised contrastive learning approaches.

In summary, while our model and data management solutions were not without limitations, they offer a path forward in the development of fast and accurate identification of cryptic biological objects in high-throughput settings.

